# AoUPRS: A Cost-Effective and Versatile PRS Calculator for the *All of Us* Program

**DOI:** 10.1101/2024.07.11.603165

**Authors:** Ahmed Khattab, Shang-Fu Chen, Nathan Wineinger, Ali Torkamani

**Affiliations:** Integrative Structural and Computational Biology, Scripps Research, La Jolla, CA, USA; Scripps Research Translational Institute, La Jolla, California, 92037, USA

## Abstract

**Background:** The All of Us (AoU) Research Program provides a comprehensive genomic dataset to accelerate health research and medical breakthroughs. Despite its potential, researchers face significant challenges, including high costs and inefficiencies associated with data extraction and analysis. AoUPRS addresses these challenges by offering a versatile and cost-effective tool for calculating polygenic risk scores (PRS), enabling both experienced and novice researchers to leverage the AoU dataset for significant genomic discoveries.

**Results:** AoUPRS is implemented in Python and utilizes the Hail framework for genomic data analysis. It offers two distinct approaches for PRS calculation: the Hail MatrixTable (MT) and the Hail Variant Dataset (VDS). The MT approach provides a dense representation of genotype data, while the VDS approach offers a sparse representation, significantly reducing computational costs. In performance evaluations, the VDS approach demonstrated a cost reduction of up to 99.51% for smaller scores and 85% for larger scores compared to the MT approach. Both approaches yielded similar predictive power, as shown by logistic regression analyses of PRS for coronary artery disease, atrial fibrillation, and type 2 diabetes. The empirical cumulative distribution functions (ECDFs) for PRS values further confirmed the consistency between the two methods.

**Conclusions:** AoUPRS is a versatile and cost-effective tool that addresses the high costs and inefficiencies associated with PRS calculations using the AoU dataset. By offering both dense and sparse data processing approaches, AoUPRS allows researchers to choose the approach best suited to their needs, facilitating genomic discoveries. The tool’s open-source availability on GitHub, coupled with detailed documentation and tutorials, ensures accessibility and ease of use for the scientific community.

## INTRODUCTION

The All of Us (AoU) Research Program, initiated by the National Institutes of Health (NIH), aims to accelerate health research and medical breakthroughs by creating a comprehensive phenotypic and genomic dataset, broadly accessible to researchers and the public. This resource includes over 245,000 short-read whole genome sequencing (srWGS) samples available in formats such as Variant Call Format (VCF), Hail MatrixTable (MT), and PLINK, all hosted on the Google Cloud Platform (GCP).

Polygenic risk score (PRS) analysis has been extensively conducted in other large, accessible databases such as the UK Biobank (Thompson *et al*. 2022), demonstrating the clinical validity of PRSs (Torkamani, Wineinger and Topol 2018). However, genomic researchers face significant challenges when executing PRS calculations with AoU data. All work must be conducted on the AoU workbench, and extracting and downloading samples to local workbenches incurs high costs. Existing tools struggle with the dataset’s scale, leading to inefficiencies and additional expenses. This contrasts with the ease and low cost of accessing health data, surveys, and other phenotypic datasets through the user-friendly AoU workbench. Consequently, there is a significant barrier, both in complexity and costs, to executing PRS calculations on AoU data.

To address these challenges, we developed AoUPRS, a versatile and cost-effective PRS calculator tool tailored for the AoU dataset. This tool is designed to facilitate both experienced and novice researchers in leveraging WGS data in AoU for genomic discoveries. Here, we present the development, implementation, and evaluation of AoUPRS, highlighting its versatility, cost-effectiveness, and potential impact on genomic research.

## METHODS

### Overview of Hail MatrixTable (MT) and Variant Dataset (VDS) Formats

Hail (https://github.com/hail-is/hail/releases/tag/0.2.130.), a scalable framework for genomic data analysis, supports two primary data formats: MatrixTable (MT) and Variant Dataset (VDS). The MT format is a dense representation where each cell (sample-variant pair) contains genotype information. This format is ideal for datasets with high variant density and allows efficient querying and manipulation of genetic data. However, it can be computationally expensive and less efficient for large-scale datasets. Accessing the Hail data in this format is currently presented in AoU tutorials for beginning researchers.

In contrast, the VDS format is a sparse representation optimized for storing large genomic datasets with many variants. It uses a more compact structure where only non-reference calls are stored, significantly reducing storage requirements and computational costs. This makes the VDS format more suitable for large-scale studies.

### AoUPRS: Approaches for PRS Calculation

AoUPRS provides two distinct approaches for calculating PRS using the AoU dataset. Each approach leverages the strengths of the respective Hail data formats to balance computational efficiency and cost-effectiveness. In both approaches, the effect allele counts are multiplied by their corresponding weights to compute weighted counts. The total PRS for each sample is then calculated by summing the weighted counts across all relevant variants. The results, including total PRS and the number of variants used, are written to an output file with an option to export all found variants contributing to the PRS for further analysis.

### Approach 1: Using Hail Dense MatrixTable (MT)

In the first approach, PRS weights are imported as a Hail Table and annotated with variant information, including effect alleles and their weights. The MT is filtered to retain only the variants present in the PRS weights table. The tool then calculates the effect allele count for each variant by comparing the reference and alternate alleles against the PRS weight table, handling different genotypic scenarios, such as homozygous reference, homozygous alternate, and heterozygous genotypes.

### Approach 2: Using Hail Sparse Variant Dataset (VDS)

In the second approach, the Variant Annotation Table (VAT) provided by the AoU Research Program is utilized. The VAT contains comprehensive annotations for all variants in the dataset, including variant identifiers, allele frequencies, and annotations across different population subgroups. Given that the VDS only stores non-reference calls, the VAT is queried to ensure that only variants present in the dataset are included in the PRS calculation, optimizing efficiency and relevance. PRS weights are imported as a Hail Table, and the VDS is filtered based on the loci specified in the PRS weights table using interval queries. The tool handles missing genotype calls by assuming they represent homozygous reference genotypes and calculates effect allele counts similarly to the MT approach.

## RESULTS

### Cost and Performance

To evaluate the performance and cost-effectiveness of AoUPRS, we compared three different PRS scores using both the Hail MT and VDS approaches. The evaluation included the cost, computational resources used, and time taken for the calculations. The results are summarized in Table 1.

**Table 1:**
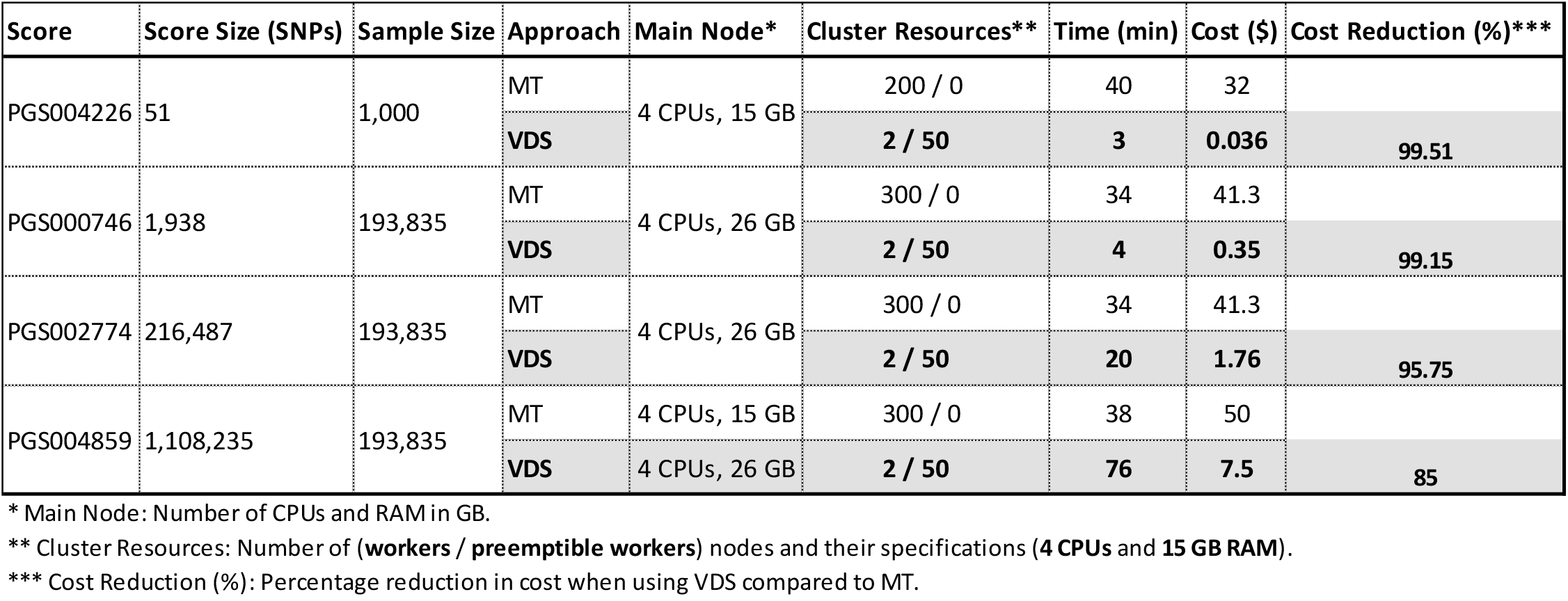
Cost, Resources, and Time for Different Scores Using Hail MT and VDS Approaches.

The results demonstrate that the VDS approach is significantly more cost-effective compared to the MT approach. For instance, the VDS approach achieved a cost reduction of 99.51% for a 51 SNP score and 85% for a score comprising 1 108 235 SNPs. The VDS format’s sparse representation minimizes computational costs, making it more suitable for large-scale studies. However, it is important to note that while the VDS approach is much cheaper, filtering intervals in the VDS tends to be slower with large weight tables, which increases the time required for calculations compared to the MT approach.

### Performance Evaluation and Predictive Power

We assessed the performance of AoUPRS by evaluating the association between the PRS for three scores from the PGS Catalog (Lambert *et al*. 2021) [PGS000746 (Gola *et al*. 2020), PGS002774 (Wong *et al*. 2022), and PGS004859 (Deutsch *et al*. 2023)] and their respective phenotypes (coronary artery disease, atrial fibrillation, and type 2 diabetes). Logistic regression analysis was conducted using the results from both the Hail MT and VDS datasets. For PGS000746, which is associated with coronary artery disease, the odds ratio (OR) was 1.090 with a 95% confidence interval (CI) of [1.077, 1.104] using MT, and 1.092 with a 95% CI of [1.079, 1.106] using VDS. For PGS002774, associated with atrial fibrillation, the OR was 1.794 with a 95% CI of [1.712, 1.879] using MT, and 1.785 with a 95% CI of [1.704, 1.870] using VDS. For PGS004859, associated with T2D, both approaches yielded an OR of 1.096 with a 95% CI of [1.072, 1.120]. These results indicate that both methods yielded similar predictive power and significance levels.

To further compare the performance of the Hail MT and VDS approaches, we generated empirical cumulative distribution functions (ECDFs) for the PRS values of the three scores plus PGS004226 (Liu *et al*. 2023) score from both approaches. The ECDF plots (see supplemental Figure 1.) illustrate that the PRS distributions generated by the Hail MT and VDS approaches are nearly identical, reinforcing the finding that both methods provide comparable results in terms of predictive power and distribution of PRS values.

## DISCUSSION

The findings from the comparison indicate that the VDS approach is more cost-effective than the MT approach, particularly for scores with a lower number of markers. Moreover, both approaches yielded virtually identical logistic regression results for all three scores, suggesting that the predictive power and significance levels are consistent across both methods. This implies that researchers can use the VDS approach to achieve cost savings without compromising the accuracy or reliability of their PRS analyses.

There are notable differences and limitations in both approaches. An Alternate Allele Count Frequency (ACAF) threshold (AF >1% and AC > 100) is imposed on the MT in AoU. This makes the MT approach susceptible to missing low-frequency variants, which can affect the completeness of the PRS, especially for larger weight tables. Conversely, the VDS approach assumes all ‘no calls’ are homozygous reference, which can lead to discrepancies. Consequently, the PRS calculated using the VDS approach are slightly different for almost all individuals compared to the MT approach, due to the impact of missing genotypes that are treated as homozygous reference in the VDS.

Given their nature and limitations, Hail MT and VDS are suited to different scenarios. Hail MT is ideal for detailed analyses requiring comprehensive genotype data and is suitable for studies with high variant density and where the computational cost is less of a concern. On the other hand, Hail VDS is best for large-scale studies where efficiency and cost-effectiveness are paramount, making it suitable for analyses involving many variants or population-level studies where sparse representation offers significant advantages. Additionally, VDS is particularly advantageous for pilot or unfunded projects, as it allows researchers to maximize their $300 free computing credit without exhausting resources as quickly as with the MT approach.

## CONCLUSION

Both Hail MT and VDS are powerful tools for PRS calculation, each with unique strengths and limitations. The choice between them should be guided by the specific requirements of the study, considering factors such as variant density, computational cost, and the importance of capturing low-frequency variants. AoUPRS demonstrates versatility and cost-effectiveness, facilitating genomic discoveries in the AoU dataset.

## ACKNOWLEDGEMENTS

We thank the All of Us team for their effort in maintaining and keeping the data available for researchers.

## DATA AVAILABILITY

AoUPRS along with a step-by-step tutorial can be found at GitHub. The repository includes the source code, documentation, and instructions for using the tool.

