## Supplemental Figure 1 for "AoUPRS: A Cost-Effective and Versatile PRS Calculator for the *All of Us* Program"

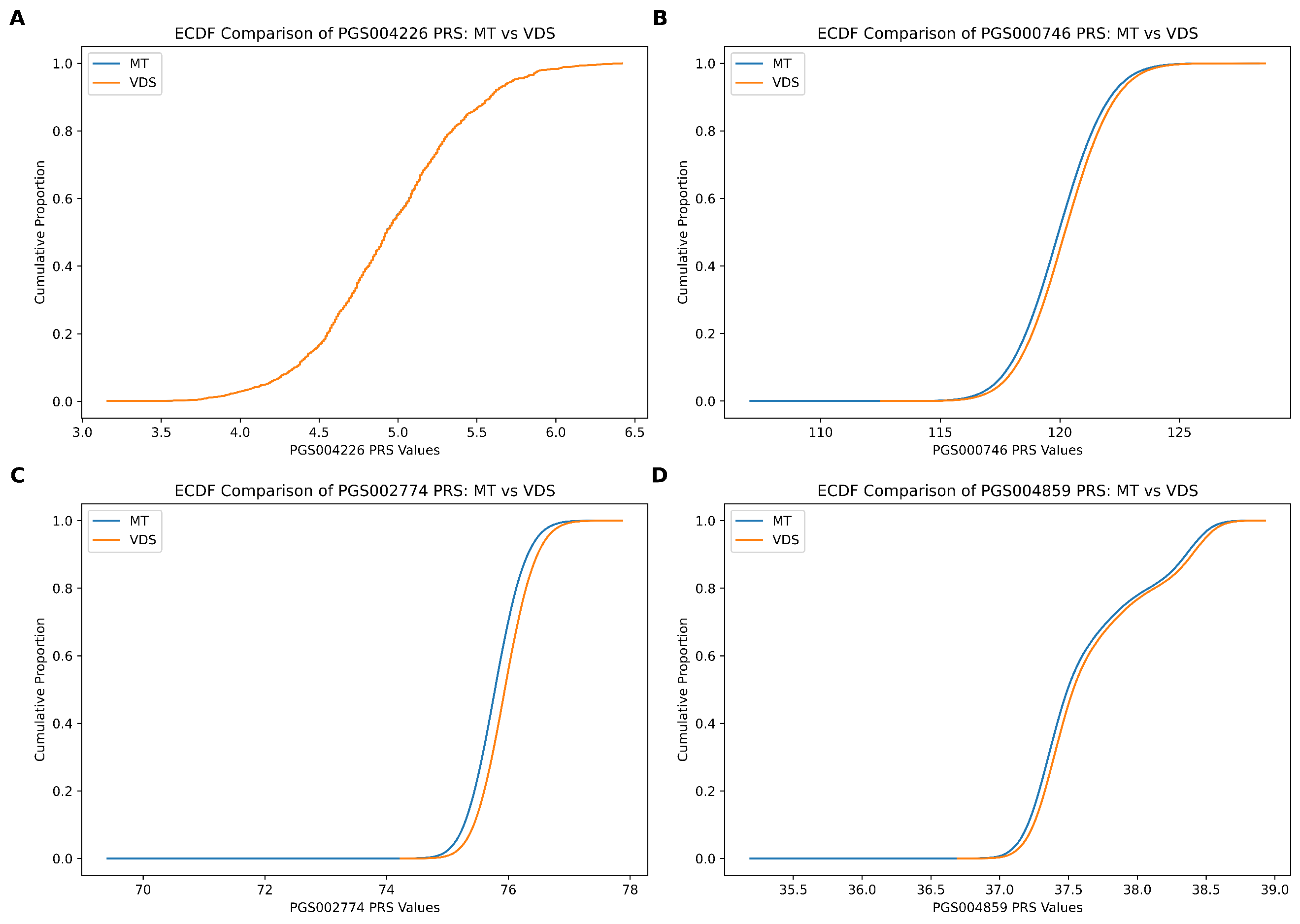

**Supplemental Figure 1:** ECDF Comparison of PRS Values Using Hail MT and VDS Approaches

Empirical cumulative distribution function (ECDF) plots comparing the polygenic risk score (PRS) values generated by Hail MatrixTable (MT) and Variant Dataset (VDS) approaches for three scores from the PGS Catalog. The plots show nearly identical distributions for each score, indicating similar predictive power and distribution of PRS values between the two methods. (A) PGS0004226 associated with type 2 diabetes, (B) PGS000746 associated with coronary artery disease, (C) PGS002774 associated with atrial fibrillation, and (D) PGS004859 associated with type 2 diabetes.
